# Neuropredictome: a data-driven predictome for cognitive, psychiatric, medical, and lifestyle factors on the brain

**DOI:** 10.1101/2020.12.07.415091

**Authors:** Syed Fahad Sultan, Lilianne R. Mujica Parodi, Steven Skiena

## Abstract

Most neuroimaging studies individually provide evidence on a narrow aspect of the human brain function, on distinct data sets that often suffer from small sample sizes. More generally, the high technical and cost demands of neuroimaging studies (combined with the statistical unreliability of neuroimaging pilot studies) may lead to observational bias, discouraging discovery of less obvious associations that nonetheless have important neurological implications. To address these problems, we built a machine-learning based classification framework, *NeuroPredictome*, optimized for the reliability and robustness of its associations. NeuroPredictome is grounded in a large-scale dataset, UK-Biobank (N=19,831), which includes resting and task functional MRI as well as structural T1-weighted and diffusion tensor imaging. Participants were assessed with respect to a comprehensive set of 5,034 phenotypes, including the physical and lifestyle factors most relevant to general medicine. Results generated by data-driven classifiers were then cross-validated, using deep-learning textual analyses, against 14,371 peer-reviewed research articles, providing an unbiased hypothesis-generator of linkages between diverse phenotypes and the brain. Our results show that neuroimaging reveals as many neurological links to physical and lifestyle factors as to cognitive factors, supporting a more integrative approach to medicine that considers disease interactions between multiple organs and systems.

## INTRODUCTION

Each year, thousands of neuroimaging studies explore the links between brain, behavior, and (primarily) psychiatric and neurological disease. Individual studies however themselves, suffer from small sample sizes and hence seldom have the statistical power to establish fully trustworthy results^1,2^. The median sample size of fMRI studies in 2015 was 28.5 subjects^3^ and the 75th percentile of sample size in cognitive neuroscience journals published between 2011 to 2014 was 28 subjects^4^. In neuroimaging, sample sizes are small due to the financial cost of scans, which can exceed $1,200 USD per data point. But working with such small sample sizes yields low statistical power and an inflated false discovery rate. As a consequence, neuroimaging has been criticized for overestimating effect sizes^5^ and concerns regarding reproducibility^6^.

Meta-analyses and literature reviews try to resolve this problem by coalescing results from a wide number of studies. Coordinate-Based Meta-Analysis (CBMA) methods^7–9^ assess the consistency of results across studies, comparing the observed spatial density of reported brain stereotactic coordinates to the null hypothesis of a uniform distribution. The latest automated CBMA methods such as Neurosynth^10^ and Neuroquery^11^ thus give excellent statistical power while enabling large scale analyses of brain-imaging studies. However, meta-analyses cannot correct for biases in the primary literature. There is a general bias in science to publish mainly significant results while experiments failing to reject the null-hypothesis are often not reported^12,13^. This in turn leads to confirmation bias by biasing future researchers to search, interpret and publish results in ways that it are in line with existing theories and hypotheses. Thus, the primary assumption required of meta-analyses, the independence of its datasets, is not always valid.

Finally, there is a problem of observational bias. An old fable has the drunkard at night searching under the streetlight for his keys, not because that is where he believes them to be, but rather because that is where the light is. Similarly, danger of purely hypothesis-based research is that it arbitrarily limits a field’s search space to the imperfect imaginations of its practitioners. Looking only at expected relationships between two variables creates a research “streetlight” that excludes other variables that may be much more important. Observational bias results from the precautions that medical research takes to minimize Type 2 error, by using highly targeted hypotheses to limit the problem of multiple comparisons. This bias is inadvertently a secondary effect of small sample sizes, because the larger the number of variables (as required for exploratory research), the greater must be the statistical power. Thus the sample size must be large in order to be able to correct for multiple comparisons without succumbing to Type 1 error.

In this paper, we present *NeuroPredictome* (NP), a hypothesis-generating tool designed to leverage recent advances in large-scale data collection and automated meta-analyses within the neuroimaging community, creating an integrative framework to minimize the problems of low reliability and reproducibility endemic to small sample sizes, confirmation bias and observational bias. Our general strategy, as shown in Figure 1, starts with a large-scale hypothesis-neutral dataset to identify linkages between brain and phenotypes, linkages that are then cross-validated against meta-analyses from the neuroimaging literature.

**Figure 1.**
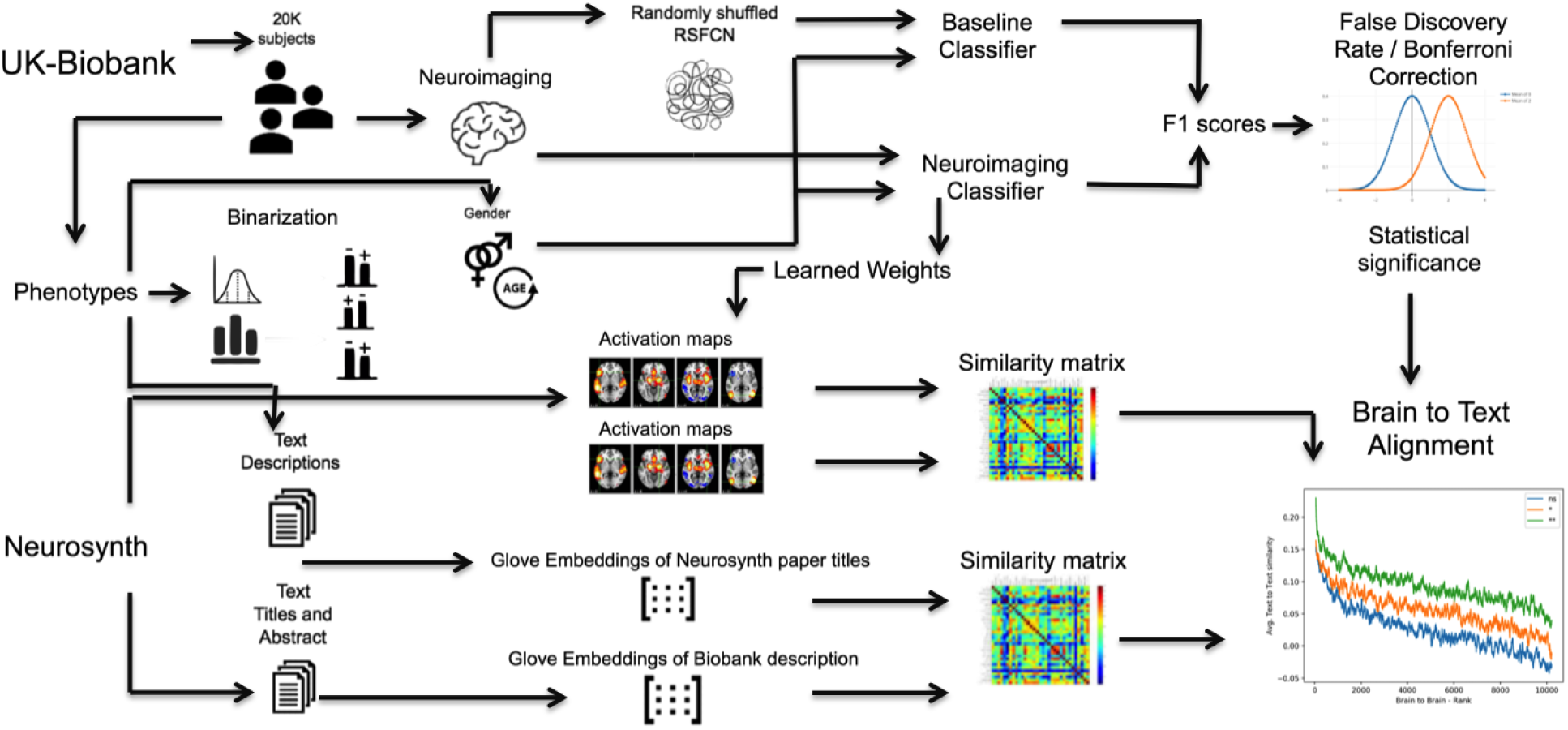
Schematic overview of NeuroPredictome. For each neuroimaging modality, brain features of 19,831 subjects, along with age and sex, are used to train 4,926 classifiers, one for each binarized-phenotype, using 20-fold cross validation. Prediction scores from each classifier is compared to that of a baseline classifier to measure statistical significance. Weights learned by each classifier is used to generate an activation map for each phenotype that is then compared to activation maps reported in Neurosynth papers. UK-Biobank phenotypes and Neurosynth papers are similarly compared using text features and we then observe how well do similarities using text features align with similarities using brain features.

For the large-scale dataset NP uses UK Biobank^14^, which to date includes neuroimaging of 19,831 individuals (as the sample size grows, NP’s pipeline will be continuously updated with new data). These neuroimaging modalities include both structural measures of neuroanatomy (T1-weighted structural brain MRI; T1) and white-matter tractography (diffusion tensor imaging; DTI), as well as functional MRI under resting and task (emotion recognition^15,16^) conditions. To link brain to phenotypes, NP binarizes 5,034 variables related to cognitive, psychiatric, medical, lifestyle, and demographic phenotypes, collected on the same individuals from whom we have neuroimaging scans. To aid in interpretability, the positive class (disease group) is always the smaller of the two classes and the negative class (control group) is always the majority class. Machine learning identifies brain features predictive of each phenotype. We model resting state fMRI as functional connectivity networks (RSFCN) and use logistic regression (LR) for classification. These together have previously been shown to exhibit predictive power comparable to more sophisticated deep learning representations and classifiers, with substantially reduced computational costs^17,18^. For each variable, we evaluate our classifier by comparing it to predictions from a baseline classifier, which uses age, gender, and randomly-shuffled features. We use 20-fold cross validation for statistical robustness, learning the model weight parameters independently for each fold. We categorize a variable as *predicted* by testing the null hypothesis that the distribution of F1-scores of our classifier and the baseline classifier are the same, and perform Bonferroni correction to account for false discoveries on account of multiple comparisons.

Those linkages that pass statistical thresholds for multiple comparisons are then cross-validated against published research studies. To do this, NP constructs a model for each BioBank phenotype that identifies brain regions differentially predictive for positive subjects. These brain maps provide a way to link phenotypes to the neuroimaging literature through the Neurosynth database, which collects brain maps associated with over 14,371 peer-reviewed published studies. Using a linguistic deep-learning measure of text-distance, NP evaluates the similarity between a phenotype description and the title of a research paper. For each phenotype, NP then orders the entire Neurosynth literature based on brain activation patterns, from most to least similar.

All associations between phenotypes and the brain that we identified can be accessed at the NP website https://neuropredictome.com. Our tool is intended to be a resource for unbiased identification of connections between brain activity and phenotypes, results from which can then be used to motivate and justify future research directions. To aid in the design of these research studies, NP provides classification scores that function as “table stakes” (baseline scores) against which researchers can compare the predictive ability of their more complex hypothesis-based models. NP also provides minimum sample sizes suggested by power analyses, thereby providing many of the benefits of neuroimaging pilot studies without their inherent unreliability and cost.

## RESULTS

### Neuropredictome identifies brain-phenotype linkages, both expected and new within neuroimaging

A summary of UK Biobank neuro-phenotype linkages is provided in Figure 2; representative variables of interest and their corresponding prediction statistics are shown in Table 1. For all variables measuring cognitive performance, subjects who scored above the median were distinguished from subjects who did worse. Many fine-grained aspects of mental health, such as depression, anxiety, mood swings, manic episodes and poor appetite were also predicted from fMRI brain activity. That psychiatric and cognitive phenotypes were linked to brain features was reassuring, but not surprising. However, one of our most striking results was that the physical and lifestyle phenotypes of greatest relevance to general medicine, while not typically assessed by neuroimaging studies, also showed brain effects. These included physical measures such as diet, Body Mass Index (BMI), and cardiovascular disorders such as angina, heart attack, strokes and blood pressure. It also included variables believed to be determinants of general well-being, such as quality of sleep, lack of social support system, experience of past trauma, satisfaction with family relationships, financial security, as well as leisure and social activities. Smoking, alcohol consumption and cannabis usage, even occasional, or in the past, was also found to alter brain activity enough for it to be strongly identifiable from resting state fMRI scans. The NP classifier was able to distinguish subjects who had never smoked cigarettes or cannabis from those who have tried it at some point, including participants who had already quit, suggesting a surprisingly long lasting footprint of addictive substances on the brain that endures long past consumption. For those that still actively consumed these substances, NP could predict the amount of consumption up to a coarse approximation. Importantly, the NP classifier passed critical “sanity checks” by failing to find relationships between brain measures and phenotypes such as bone fractures and month of birth. Likewise, brain measures did not predict ethnicity.

**Table 1.** Neuropredictome identifies brain-phenotype linkages, across different phenotype categories, that have gone ignored in neuroimaging along with some linkages that were to be expected.. A linkage is considered strongly statistically significant if p-value <0.001 (**) and significant if p-value <0.01 (*) based on rejection of the null-hypothesis of association between F1-score distributions, from different folds of cross-validation, using Resting-State Functional Connectivity Network (RSFCN) and F1-score distribution from baseline classifier (Base) after Bonferroni correction.

| Category | Variable | +ve pct | F1 (Base) | F1 (RFCN) | p-value | sig* |
| --- | --- | --- | --- | --- | --- | --- |
| Health and medical history | Fractured/broken bones in last 5 years | 0.083 | 0 | 0.00417 | 0.33 |  |
| Health and medical history | Fracture resulting from simple fall rest (+ve) vs Yes | 0.429 | 0.496 | 0.501 | 0.405 |  |
| Health and medical history | Fractured bone site(s) Ankle (+ve) vs rest | 0.0938 | 0.0178 | 0.0538 | 0.000241 |  |
| Health and medical history | Fractured bone site(s) Wrist (+ve) vs rest | 0.181 | 0.0528 | 0.0678 | 0.0167 |  |
| Lifestyle and environment | Major dietary changes in the last 5 years rest (+ve) vs No | 0.36 | 0.03 | 0.09 | 4.13e-18 | ** |
| Lifestyle and environment | Beef intake > Less than once a week | 0.394 | 0.0951 | 0.151 | 7.87e-16 | ** |
| Lifestyle and environment | Coffee intake > 75th percentile | 0.19 | 0 | 0.000832 | 0.00211 |  |
| Health and medical history | Diabetes diagnosed by doctor Yes (+ve) vs rest | 0.0499 | 0 | 0.000375 | 0.33 |  |
| Lifestyle and environment | Number of cigarettes currently smoked daily < 50th percentile | 0.469 | 0.469 | 0.523 | 1.05e-06 | ** |
| Lifestyle and environment | Smoking status Previous (+ve) vs rest | 0.336 | 0.102 | 0.132 | 1.37e-10 | ** |
| Lifestyle and environment | Ever taken cannabis Yes (+ve) vs No | 0.215 | 0.0796 | 0.136 | 2.35e-12 | ** |
| Lifestyle and environment | Frequency of drinking alcohol ≤ Monthly or less | 0.197 | 0.00114 | 0.0186 | 2.22e-09 | ** |
| Lifestyle and environment | Frequency of drinking alcohol > 2 to 3 times a week | 0.303 | 0.0701 | 0.123 | 8.97e-14 | ** |
| Cognitive Function | Fluid intelligence score < 50th percentile | 0.455 | 0.345 | 0.485 | 1.55e-23 | ** |
| Cognitive Function | Matrix Pattern Completion < 50th pctl. | 0.372 | 0.279 | 0.378 | 2.33e-17 | ** |
| Cognitive Function | Numeric Memory < 50th percentile | 0.338 | 0.0987 | 0.194 | 5.35e-17 | ** |
| Cognitive Function | Reaction Time < 50th pctl. | 0.492 | 0.603 | 0.614 | 2.39e-06 | * |
| Cognitive Function | Trail making < 50th percentile | 0.493 | 0.611 | 0.626 | 8.52e-07 | ** |
| Cognitive Function | Tower rearranging < 50th percentile | 0.405 | 0.358 | 0.411 | 6.18e-11 | ** |
| Mental Health | Seen doctor (GP) for nerves, anxiety, tension or depression | 0.321 | 0.104 | 0.138 | 2.05e-11 | ** |
| Mental Health | Bipolar and major depression status > No Bipolar or Depression | 0.291 | 0.114 | 0.18 | 7.23e-11 | ** |
| Mental Health | Manic/hyper symptoms None of the above (+ve) vs rest | 0.43 | 0.391 | 0.429 | 9.18e-10 | ** |
| Mental Health | Family relationship satisfaction > Very happy | 0.322 | 0.0177 | 0.0415 | 9.69e-11 | ** |
| Mental Health | Financial situation satisfaction > Very happy | 0.408 | 0.301 | 0.325 | 4.83e-10 | ** |
| Mental Health | Leisure/social activities None of the above (+ve) vs rest | 0.245 | 8e-05 | 0.01 | 2.9e-11 | ** |
| Mental Health | Victim of sexual assault > Never | 0.152 | 0.000532 | 0.00596 | 5.78e-06 | * |
| Mental Health | Physically abused by family as a child > Never true | 0.199 | 0.000269 | 0.00468 | 5.61e-07 | ** |
| Physical measures | Body mass index (BMI) > 75th percentile | 0.248 | 0.000736 | 0.145 | 1.5e-23 | ** |
| Physical measures | Diastolic blood pressure, manual reading > 75th percentile | 0.229 | 0.0925 | 0.141 | 1.38e-07 | ** |
| Physical measures | IPAQ activity group > moderate | 0.397 | 0.103 | 0.172 | 6.12e-17 | ** |
| Lifestyle and environment | Sleeplessness / insomnia ≤ Never rarely | 0.221 | 0.00345 | 0.011 | 2.5e-07 | ** |
| Lifestyle and environment | Daytime dozing / sleeping (narcolepsy) Sometimes (+ve) vs rest | 0.205 | 0.00168 | 0.0176 | 7.54e-10 | ** |
| Lifestyle and environment | Sleep duration < 25th percentile | 0.244 | 0.000247 | 0.012 | 8.74e-10 | ** |
| Sociodemographics | Average total household income before tax ≤ 18,000 | 0.213 | 0.0138 | 0.0561 | 9.99e-15 | ** |
| Sociodemographics | Average total household income before tax > 31,000 | 0.254 | 0.256 | 0.309 | 2.89e-13 | ** |
| Sociodemographics | Ethnic background African (+ve) vs rest | 0.00199 | 0 | 0.0161 | 0.109 |  |
| Sociodemographics | Ethnic background Chinese (+ve) vs rest | 0.00277 | 0 | 0.031 | 0.0153 |  |
| Sociodemographics | Ethnic background Indian (+ve) vs rest | 0.00654 | 0 | 0.0295 | 0.00475 |  |
| Sociodemographics | Ethnic background rest (+ve) vs British | 0.0744 | 0 | 0.0043 | 0.00122 |  |
|  | UK Biobank assessment center Newcastle (+ve) vs Cheadle | 0.156 | 0 | 0.246 | 1.2e-24 | ** |

**Table 2.** Activation Maps learned by Neuropredictome for a wide variety of phenotypes strongly align with relevant Neurosynth papers. A curated list is given of UK-Biobank phenotypes and their corresponding top ranked Neurosynth paper with respect to similarity in reported brain activations and those identified by Neuropredictome. Corresponding text similarity between description of phenotype and title and abstract of top-ranked paper is also given.

| ID | Biobank variable | pval | Text similarity | Neurosynth paper title | PubMed ID |
| --- | --- | --- | --- | --- | --- |
| 20116 | Smoking status > Never | 2.15e-16 | 0.21 | Nicotine withdrawal modulates frontal brain function during an affective Stroop task | 21989805 |
| 20414 | Frequency of drinking alcohol > 2 to 3 times a week | 8.97e-14 | 0.2 | Influence of cue exposure on inhibitory control and brain activation in patients with alcohol dependence. | 22557953 |
| 5699 | Fluid intelligence: arithmetic sequence recognition | 5.52E-17 | 0.13 | The interaction of process and domain in prefrontal cortex during inductive reasoning | 25498406 |
| 20240 | Numeric Memory - Maximum digits remembered correctly < 50th percentile | 5.35e-17 | 0.17 | Cross-modal processing in auditory and visual working memory | 16154767 |
| 6772 | Trail making - Interval between previous point and current one in numeric path >75 percentile | 4.26E-14 | 0.22 | The neural basis for establishing a focal point in pure coordination games. | 22009019 |
| 2090 | Seen doctor (GP) for nerves, anxiety, tension or depression | 4.26E-14 | 0.36 | Regional homogeneity associated with overgeneral autobiographical memory of first-episode treatment-naïve patients with major depressive disorder in the orbitofrontal cortex | 27923192 |
| 20425 | Anxiety - Ever worried more than most people would in similar situation | 2.01E-12 | 0.22 | Metabolic and functional connectivity changes in mal de debarquement syndrome | 23209584 |
| 20126 | Bipolar II Disorder (+ve) vs rest | 2.87E-11 | 0.36 | Possible structural abnormality of the brainstem in unipolar depressive illness | 16227541 |
| 21002 | Weight > 75% percentile | 6.97E-24 | 0.24 | Widespread reward-system activation in obese women in response to pictures of high-calorie foods. | 18413289 |
| 1220 | Daytime dozing / sleeping (narcolepsy) | 3.64E-13 | 0.13 | Hippocampal activity mediates the relationship between circadian activity rhythms and memory in older adults. | 26205911 |

**Figure 2.**
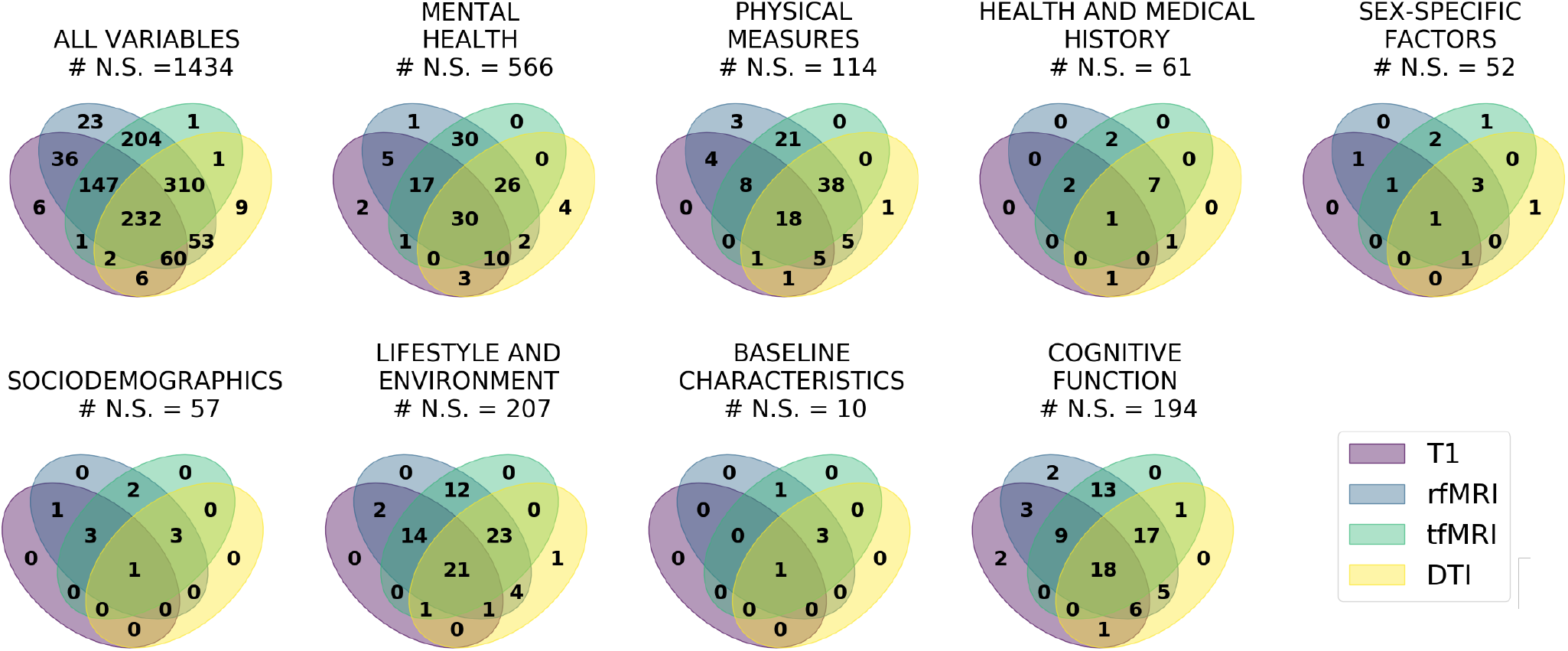
Neuropredictome identifies new brain-phenotype linkages, along with some expected ones. Physical and lifestyle phenotypes, typically not looked at in light of neuroimaging, showed strong brain effects. Across neuroimaging modalities, resting state functional connectivity (rfMRI) broadly yields the best prediction scores and predicts a larger number of phenotypes compared to structural DTI, T1-weighted or task fMRI.

Comparing across neuroimaging modalities, our results show Resting-state fMRI to broadly yield the best prediction scores and to predict the largest number of phenotypes as compared to structural DTI, T1-weighted or task fMRI. In line with previous findings^19^, our results show that resting functional connectivity lends *additive* prediction power to structural neuroimaging and is a more generally informative fingerprint of the subject as compared to task networks^20^.

Also of note for multi-site studies was the profound effect of imaging site. This was despite the fact that in order to maximize data compatibility across the different imaging centers, identical scanners were used with no major software or hardware updates throughout the study, with identical acquisition parameters, as well as identical post-processing pipelines. These results are in line with recent work by our group showing the influence of scanner-specific artifacts on neuroimaging data^21^.

### Brain maps learned by our classifier align with results from 14,371 published papers

As shown by Figure 3, Neuropredictome’s classifiers from Biobank linkages were strongly supported by Neurosynth meta-analyses of the neuroimaging literature. For each category, if our brain-to-brain similarity matrix indicated a high similarity, yielding a high rank (with the highest similarity or rank represented by the smallest difference between them, zero), then the corresponding text-to-text similarity score was also high. Furthermore, for the variables for which our prediction was most accurate (**), we obtain a sharper slope than for the variables for which we did moderately well (*), which in turn has a sharper slope than the set of variables for which our prediction results were uninformative (n.s.). This separation of the three lines, even at the tail end towards the right end of the figure, is indicative of the meaningfulness that is to be attributed to the weights obtained using the learning process. For variables that we were not able to predict, the weights for the brain regions were either meaningless and thus the resultant brain activation map was also a null map, more or less uniformly distributed. In case of variables that we could not predict, it is also important to point out that the highest weights also often went to age and/or sex variables, which itself is informative.

**Figure 3.**
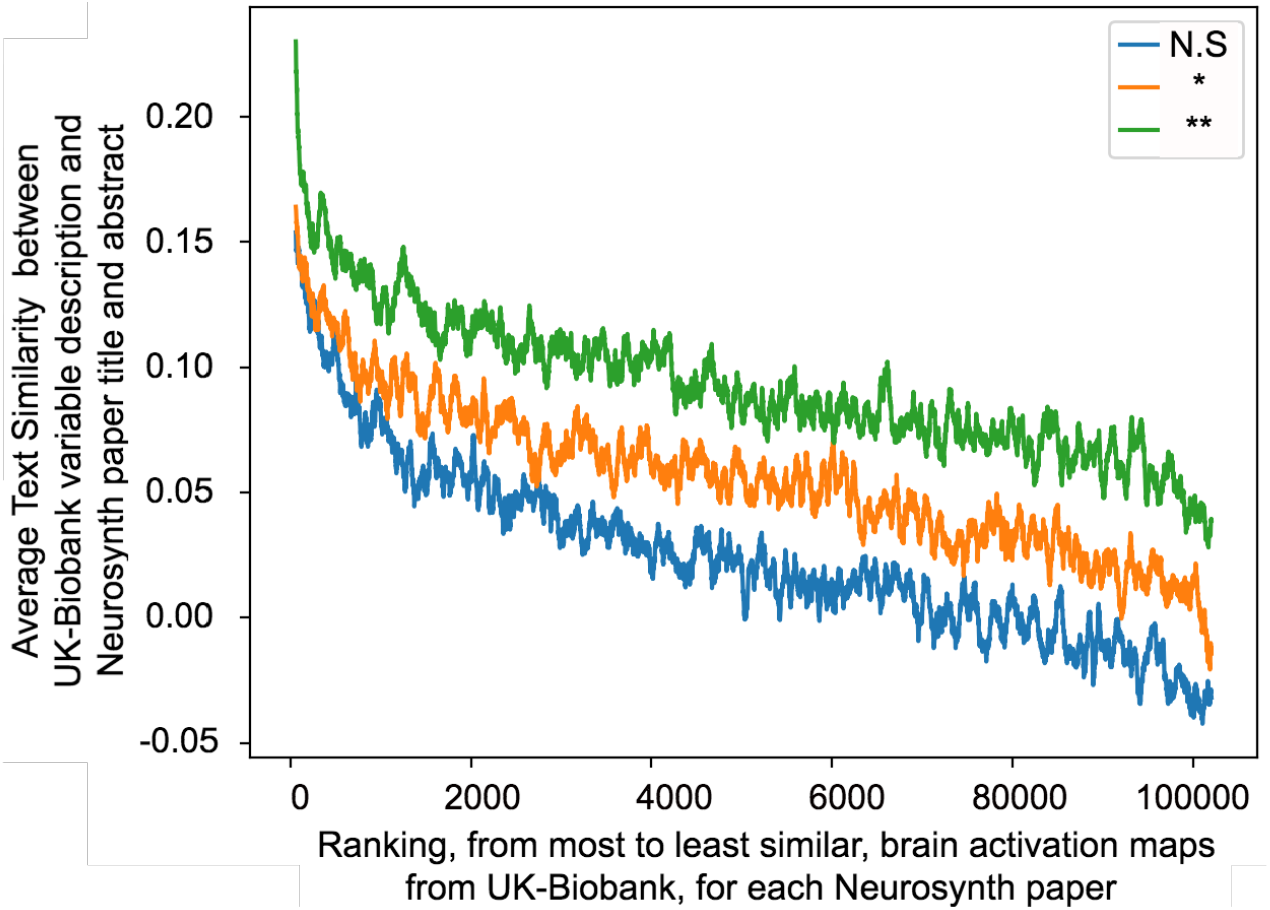
Neuropredictome results strongly align with results reported in 14,371 Neurosynth papers. When Neuropredictome maps a Neurosynth paper to a UK-Biobank phenotype based on similarity in brain activity, then the phenotype-paper pair are also similar in terms of textual descriptions. The more confident Neuropredictome is about its predictions, the stronger the alignment. Most significant phenotypes (**) *r* = 0.25; moderately significant *r* = 0.18 (*); for non-significant (N.S) phenotypes *r* = 0.09.

## DISCUSSION

In this paper, we present NeuroPredictome (NP), a machine learning tool designed to avoid observational bias by allowing large-scale hypothesis-neutral data to guide us in identifying new relationships between brain-based features and phenotypes. Some of these phenotypes (e.g., psychiatric and cognitive variables) are already of significant interest to the neuroimaging community. Thus, in spite of recent and well-warranted criticism regarding reproducibility and reliability concerns within the field, our findings provide reassurance that neuroimaging remains a methodology of non-trivial detection sensitivity in understanding the brain and its relationship to brain-based disease.

More provocatively, however, our results suggest greater integration between the brain and disorders of other organs and systems than is typically assumed. Just as endocrine disorders such as Graves Disease and diabetes can produce symptoms of anxiety and cognitive decline, respectively, in disease processes the lines between specializations are often less segregated than blurred. Thus, by providing hypothesis-neutral estimates of modality-specific relationships and the statistical power required to detect them, NP can provide an alternative to the problem of expensive pilot studies, plagued by low detection sensitivity and unreliable effect size estimates, in motivating future work in bold and unexpected new directions.

One potential limitation of this work is its reliance on a particular dataset that, in spite of breadth, still (of necessity) had to make choices about which phenotype variables to include or exclude. To address this, we plan to expand the NP framework to incorporate additional large-scale (N>1,000) neuroimaging datasets as they become available, with the most recent updates posted on our website https://neuropredictome.com. A second potential limitation is our dependence on one particular model and classifier of brain activity, as one might wonder if different models and classifiers could yield different results. However, to date our model and classifier represents the state of the art in terms of its applications to neuroimaging data, as recent work^17^ shows resting state functional networks and kernel ridge regression classifier to produce results comparable to more sophisticated deep learning based models on the same Biobank UK data set.

Building on that methodological work, we expand its scope by studying a comprehensive set of conditions against brain activity, as well as by cross-validating our findings with meta-analyses. Our integration of UK Biobank classifiers with NeuroSynth provides important reassurance that, whatever the limitations present in individual neuroimaging studies, the neuroimaging field as a whole shows convergence with a dataset large enough to be considered ground truth.

## Acknowledgments

The research described in this paper was funded by the W. M. Keck Foundation (L.R.M.-P.) and the White House Brain Research Through Advancing Innovative Technologies (BRAIN) Initiative (NSFNCS-FR 1926781 to L.R.M.-P.), and by NSF grants IIS-1926751, IIS-1927227, and IIS-1546113 and the New York Empire Innovation Program (to S.S.).

## Data Availability

The neuroimaging and phenotypic data used in this work was obtained from UK-Biobank under Data Access Application 37462 and is available upon application (http://www.ukbiobank.ac.uk/register-apply/).

## Code Availability

All source code is available from the corresponding author upon reasonable request.

## METHODOLOGY

### UK-Biobank Dataset

The UK Biobank^22^ is a prospective epidemiological study that recruited 500,000 adults (age 40-69) between 2006-2010, with 100,000 of these 500,000 participants brought back for multimodal imaging by 2022^23^. Here, we considered an initial release of around 19,831 subjects with both structural MRI and rs-fMRI data. Subjects recruited in the UK-Biobank study are predominantly of middle and older age. In addition to brain imaging data for these subjects, the study also collects data from extensive questionnaires, physical and cognitive measures, and biological samples (including genotyping). All participants consented to allowing access to their full health records from the UK National Health Service, enabling researchers to relate phenotypic measures to long-term health outcomes. This is particularly powerful as a result of the combination of the number of subjects and the breadth of linked data.

Participants ages range from 40 to 69 years of age at baseline recruitment; this aims to balance the goals of characterizing subjects before disease onset against the delay before health outcomes accumulate. The cohort is particularly appropriate for the study of age-associated pathology. For example, in the imaged cohort, 1,800 participants are expected to develop Alzheimer’s disease by 2022, rising to 6,000 by 2027 (diabetes: 8,000 rising to 14,000; stroke: 1,800 to 4,000; Parkinson’s: 1,200 to 2,800). Along with body and cardiac imaging, genetics, lifestyle measures, biological phenotyping and health records, this imaging is expected to enable discovery of imaging markers for a broad range of diseases at their earliest stages, as well as provide unique insight into disease mechanisms.

Both structural MRI and rs-fMRI were acquired on harmonized Siemens 3T Skyra scanners at four UK Biobank imaging centres (Cheadle, Manchester, Newcastle, and Reading). The structural MRI was 1.0mm isotropic. The rs-fMRI was 2.4mm isotropic with TR of 0.735s and 490 frames per run (6 min). Each subject had one rs-fMRI run. A number of behavioral measures were also collected by the UK Biobank.

The neuroimaging data available to us from UK-Biobank was resting state fMRI of N=19,831 subjects, with over 5034 associated auxillary labels for each subject, collected from extensive questionnaires, physical and cognitive measures, biological samples (including genotyping) and health records from the UK National Health Service.

### Binarization

Of the 5034 variables in the UK-Biobank dataset, an overwhelming majority of them were continuous or multi-class categorical variables. In this study, we restricted ourselves to the problem of binary classification in case of each condition. Even though, classifiers such as logistic regression are well suited for prediction of non-binary labels, without binarization, for certain variables we would have to carry out regression, as opposed to other classification variables. We feel that the results would have been harder to interpret. This binarization step ensures consistency across prediction of all variables, categorical or numerical.

Therefore in our experiments, each one of the available labels was converted into multiple binarized variants. For each categorical variable, we performed converted it into a binary variant for each value that variable takes on vs. rest. Additionally, since not all labels were available for all the subjects, an *available vs. not-available* variant was also created for each categorical variable. Most continuous labels available to us from the UK-Biobank data set exhibited Gaussian or Power law distributions. Each of these converted into 3 binarized variants, with respect to the three quartiles: i) <= 25th quartile vs. rest ii) <=50th quartile vs. rest and iii) <=75th quartile vs. rest. Once the binarized variants of all the labels were computed, variants that were available for at least 1000 subjects and for which the minority class was at least one percent of all the subjects were retained and the remaining were dropped. After creating the binarized variants and applying the aforementioned criterion for minimal availability and distributions, we finally end up with 4926 phenotype and variants.

### Resting State Functional Connectivity (RSFC)

Resting State Functional Connectivity (RSFC) measures the congruity between different regions of the brain while the participants are lying at rest without any explicit task. This congruity is measured from the resting-state functional magnetic resonance (rs-fMRI) image signals for each brain region. For a given parcellation scheme, the parcels are used as regions of interest (ROI) such that a whole brain (or cortical) RSFC matrix can be computed for each participant. Each entry of the RSFC matrix corresponds to the strength of the functional connectivity between two brain regions.

### FC-based prediction setup

RSFC data of each particpant was summarized as an *N* × *N* matrix, where *N* is the number of brain ROIs. Each entry in the RSFC matrix represented the strength of the functional connectivity between the corresponding pair of ROIs. The entries of the RSFC matrix were used as features to predict behavioral and demographic measures in individual participants.

### Task functional Magnetic Resonance Imaging (tfMRI)

Task functional MRI (“tfMRI”) uses the same measurement technique as resting-state fMRI, while the subject performs a particular task or experiences a sensory stimulus. The task used is the Hariri faces/shapes “emotion” task^15,16^, as implemented in the HCP, but with shorter overall duration and hence fewer total stimulus block repeats. The participants are presented with blocks of trials and asked to decide either which of two faces presented on the bottom of the screen match the face at the top of the screen, or which of two shapes presented at the bottom of the screen match the shape at the top of the screen. The faces have either angry or fearful expressions. This task was chosen to engage a range of sensory, motor and cognitive systems. The features from task fMRI scans used for classification were also functional connectivity weights, comparable to those computed for resting state fMRI.

### Diffusion Tensor Imaging (DTI)

Diffusion-weighted imaging measures the ability of water molecules to move within their local tissue environment. Water diffusion is measured along a range of orientations, providing two types of useful information. Local (voxel-wise) estimates of diffusion properties reflect the integrity of microstructural tissue compartments (e.g., diffusion tensor and NODDI measures). Long-range estimates based on tract-tracing (tractography) reflect structural connectivity between pairs of brain regions. For the two diffusion-weighted shells, 50 diffusion-encoding directions were acquired (with all 100 directions being distinct). Acquisition phase-encoding direction for the dMRI data is AP (Anterior-to-Posterior). In order to generate appropriate fieldmaps to carry out geometric distortion correction for EPI images. Fractional anisotropy, tensor mode and mean diffusivity are obtained using DTI fitting tool DTIFIT^24^ on b = 1000 shell (50 directions).

The full two-shell dMRI data is fed into NODDI (Neurite Orientation Dispersion and Density Imaging)^25^ modelling. This aims to generate meaningful voxelwise microstructural parameters, including ICVF (intra-cellular volume fraction - an index of white matter neurite density), ISOVF (isotropic or free water volume fraction) and ODI (orientation dispersion index, a measure of within-voxel tract disorganisation).

The DTI Fractional Anisotropy (FA) image is aligned onto a standard space for white matter skeletons using TBSS^26^. The resulting standard-space warp is applied to all other DTI/NODDI output maps. For each of the DTI/NODDI maps, the skeletonised images are averaged within a set of 48 standard-space tract masks defined by Susumu Mori^27^.

Within-voxel modelling of multi-fibre tract orientation structure is carried out using the bedpostx tool on the preprocessed dMRI data. 27 major tracts using standard-space start/stop ROI masks defined by AutoPtx^28^ are obtained. In addition, probabilistic tractography with crossing fibre modelling is performed using probtrackx^29^. As features to the classifier, we use for each tract, and for each DTI/NODDI output image type, the weighted-mean value of the DTI/NODDI parameter within the tract, the weight being the tractography probabilistic output.

### Structural T1-weighted imaging

T1-weighted imaging is a structural technique with high-resolution depiction of brain anatomy, having strong contrast between grey and white matter, reflecting differences in the interaction of water with surrounding tissues (tissue T1 relaxation times). This modality provides derived fields primarily relating to volumes of brain tissues and structures. It is also critical for calculations of cross-subject and cross-modality alignments, needed in order to process all other brain modalities. Acquisition details of the data avaiable to us are: 1 mm isotropic resolution using a 3D MPRAGE acquisition. The superior-inferior field-of-view is large (256 mm), at little cost, in order to include reasonable amounts of neck/mouth.

Next, tissue-type segmentation is applied using FAST^30^. This estimates discrete and probabilistic segmentations for CSF (cerebrospinal fluid), grey matter and white matter. As part of the segmentation, intensity bias is estimated, and so this step is also used to generate a fully bias-field-corrected version of the brain-extracted T1.

The external surface of the skull is estimated from the T1, and used to normalise brain tissue volumes for head size (compared with the MNI152 template) and volumes of different tissue types and total brain volume, both unnormalised and normalised for head size, are then generated, all using SIENAX^31^.

Subcortical structures were modelled using FIRST^32^. The shape and volume outputs for 15 subcortical structures are generated and stored. From the T1 structural image, several global volume measures are used as features to the classifier, both normalised for overall head size as well as non-normalised: total brain (grey + white matter) volume; total white matter volume, total grey matter volume, ventricular (non-peripheral) CSF (cerebrospinal fluid) volume; peripheral cortical grey matter volume. Several subcortical structures’ volumes from FIRST segmentation are also used, in general with separate features for left and right hemispheres. Total volume of grey matter within 139 grey matter ROIs and total volume of white matter hyperintensities are also used.

Number of features and model complexities for the four modalities are not strictly comparable. Unfortunately, this is a limitation of comparing different modalities. It is simply not feasible to ensure parity of model complexity across different modalities because each requires different standard preprocessing, and different parcellations and atlases. The model complexity across task and resting state fMRI was the same however. We addressed concerns regarding greater overfitting of a more expressible model by using 20-fold cross-validation for each modality and each phenotype.

### Confound modeling

Confounds can be significant in addressing problems of unexplained variance and spurious correlations. In UK-Biobank, the resulting high statistical power from the large number of subjects, raises the question of effect of confounds. Alfaro et al.^33^ describe a set of possible confounds in UK-Biobank that effect the data and the spurious correlations that may arise if they are not controlled. Alfaro et al.^33^ show that imaging can be influenced by blood pressure, bone density, height and weight but are also likely to partially reflect physiological characteristics of interest while also discussing Berkson’s paradox of deconfounding. Age and sex in particular are confirmed as one of the most important confounds that mediate large amounts of between-subject variance. Some of the imaging-derived confounds however were found to be obvious proxies for certain phenotypes e.g. head size is an obvious proxy for height.

Non-imaging phenotypes, the ones Neuropredictome sets out to predict from neuroimaging, also exhibit correlations amongst each other. However, most of these inter-correlated variables are within phenotype category boundaries. As a result of this, the presentation of prediction results in Figure 2 is with respect to categories of phenotypes and Table 1 presents a synthesis and hand curated list of results that avoid such inter-correlated variables.

### Normalizing for Age and Sex

Preliminary results of our experiments, in agreement with existing literature^17,18,33,34^, showed that fMRI scans can reveal the subject’s age and sex with a high degree of accuracy. Thus naturally, the variables on which we were getting the best scores were proxies for Age and sex. These variables comprised most of the labels under the categories *Sex-specific factors* and *Socio-demographics*. Given the fact that age and sex can be a strong confounding factor in most, if not all, more interesting variables, we corrected for age and sex by providing them as features to both our RSFCN classifier and the Baseline classifier. Age and sex as set of confounds have been used in many brain imaging studies. A detailed discussion on studies for which these confounds may be useful can be found in Barnes et al.^34^.

### Training, validation and testing

The large UK Biobank dataset allowed us the luxury of splitting the 19,831 subejcts into training (N=14875) and test(N=4926) sets. Care was taken so that the distribution of various attributes were similar across training, validation and test sets. Hyperparameters were tuned using the training and validation sets. The test set was only utilized to evaluate the final prediction performance. The hyperparameter *λ* was tuned using the validation set based on a grid search over the hyper parameter.

### Baseline

As baseline model, we used a Logistic Regression classifier with features age and sex. In order to have parity in terms of the number of features in our model and that of the baseline, we provided the baseline model with randomly shuffled fMRI features. In other words, the fMRI features available to the baseline model did not have the original subject to fMRI features thus encoding useless information yet preserving the overall properties of the fMRI features.

### 20-fold Cross Validation

In order to evaluate how impressive the results of our learned classifier were from random chance, we set out to compute, record and report statistical significance scores for our models. We created 20 different splits for subjects into 20 different sets of training and testing subjects. The splitting was done with the train to test ratio of 75:25. For each split, we trained age-sex matched classifiers on the training subjects and tested on the test subjects. For each one, we computed f1 scores from prediction of our RSFCN classifier and Naive Baseline model against the true labels. For all classifications, simple Logistic Regression classifiers were used. This gave us two distributions of f1-scores, each with 20 constituent data points, one for each of the 20 folds, for both, our model and the baseline.

### F1-scores

Generally, accuracy is a standard metric for judging the quality of the classifiers, when classes are balanced. However, many of the phenotypes in UK-Biobank are quite rare, creating an imbalanced classification problem. For these, majority classifiers, which always pick the largest class, can achieve trivially high accuracy. The F-1 metric that we use is the standard approach for judging classifiers in non-balanced cases.

It is calculated from the precision and recall of the test, where the precision is the number of correctly identified positive results divided by the number of all positive results, including those not identified correctly, and the recall is the number of correctly identified positive results divided by the number of all samples that should have been identified as positive.

The F1 score is the harmonic mean of the precision and recall. The highest possible value of F1 is 1, indicating perfect precision and recall, and the lowest possible value is 0, if either the precision or the recall is zero.

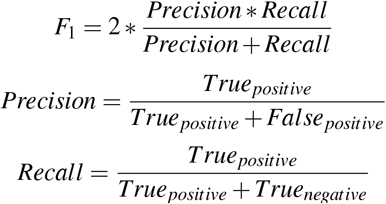

### Computing Statistical significance

The F1-score distributions were inherently Gaussian, and so to measure how distinct the distribution of our model was compared to the baseline, we performed simple t-tests between distribution of our model and the mean of the baseline. This yielded a p-value for the null hypothesis that the baseline model’s mean prediction was sampled from the distribution of prediction scores of our model. We use this p-value and its associated t-test statistic as the two primary measures of quality of our models.

### Bonferroni correction

Given the large number of labels, it is important to keep in mind that for some of the labels, the classification scores will be high just by chance. This problem is generally called the multiple comparisons problem. Therefore, comparing scores from our classifier to those of a random classifier and a simple majority class classifier - ones that did not really learned anything from the data, is important. We used Bonferroni correction to control for false discovery rates and to evaluate the quality or reliability of the accuracy score. The p-value scores in the results presented were after applying the Bonferroni correction. Please note that Bonferroni correction was performed across phenotypes for each neuroimaging modality, and not across modalities.

### RSFCN

We utilized the 55×55 RSFC (Pearson’s correlations) matrices.^23,35^. The 55 ROIs were obtained from a 100-component whole-brain spatial-ICA^36^ of which 45 components were considered to be artifactual^23^.

### Logistic regression

Logistic regression^37^ is a classical non-parametric machine learning algorithm. Let *y* be the behavioral measure and *c* be the RSFC matrix of a test subject. Let *y*_*i*_ be the behavioral measure and *c*_*i*_ be the RSFC marix of the *i*-th training subject. Roughly speaking, logistic regression will predict the test subject’s behavioral measure to be the weighted

average of the behavioral measures of all the training subjects: *y* ≈ ∑_*i*∈*trainingset*_ *Similarity*(*c*_*i*_, *c*)*y*_*i*_ where *Similarity*(*c*_*i*_, *c*) is the similarity between the RSFC matrix of the test subject and the *i*-th training subject. Here, we set *Similarity*(*c*_*i*_, *c*) to the Pearson’s correlation between the lower triangular entries of matrices *c*_*i*_ and *c*, which is effectively a linear regression. In practice, an *l*_2_ regularization term is needed to avoid overfitting (i.e. kernel ridge regression). The level of *l*_2_ regularization is controlled by the hyperparameter *λ*.

## Evaluation against Neurosynth

Neurosynth^10^ is an automatically generated repository of activation maps that associates not only an activation map with 14,371 neuroimaging papers, but also aggregates information from these activation maps with respect to coordinates in the standard MNI space and the general common terms occurring in the title and abstract of these papers. Neurosynth represents the state-of-the-art in the neuroimaging community’s understanding of organization of human brain mapping relating regions in the brain to a wide array of behavior and phenotypes. Given the availability of large sets of phenotypes in Biobank, we set to find out how much of what we learned from Biobank was in agreement with the published literature generated by the neuroimaging community over decades of research. We did this by aligning knowledge gained from Biobank with knowledge learned by the neuroimaging community as captured by Neurosynth. Details of the alignment process is given below.

### Neurosynth Brain to Biobank Brain mapping

Each of our 4,926 classifiers, trained against a particular phenotype or its variant, learns weights on the given features during the training phase. These weights are used to compute an activation map for the respective phenotype variant. Since the features we use in the classifier are Resting State Functional Connectivity Networks, the learned weights are with respect to connections (edge in a graph) between pairs of regions of interest (ROI). This is carried out in the following three steps. First, for each classifier, we average learned weights from each fold of the 20-fold cross validation. Second, we map a learned weight for each edge in the connectivity graph matrix to the corresponding pairs of ROIs (nodes of the connectivity graph). Now, since a) the ROIs we use in our experiments are ICA components, these ROIs are not necessarily disjoint and often have regions overlapping with each other and b) the Neurosynth activation maps that we seek to compare against are in voxel space, so as the third final step we map ROI weights to voxel weights. Since ROIs are essentially sets of voxels, we assign to each voxel the weight of its respective ROI. In case of voxels that fall into more than one ROI, we assign it the average weight of all its corresponding ROIs.

Once we have an activation map for each of our 4926 phenotype variants and an activation map for each of the 14,371 papers in Neurosynth, we flatten all maps into vectors. These vectors have 228,453 dimensions, equal to the number of grey matter voxels in the standard MNI coordinate space. We then compute a pairwise similarity matrix (4926,14371) between UK-Biobank phenotypes and Neurosynth paper embeddings, using Pearson correlation coefficient as the similarity metric.

### Neurosynth text to Biobank text mapping

In order to have a quantitative measure of quality for the brain to brain similarity matrix from section above, we compared text descriptions of the 4,926 UK-Biobank phenotypes with the text from abstract and title of the 14,371 Neurosynth papers. We do so by treating the Biobank variable descriptions of the phenotypes as text documents. Similarly, we concatenate the title and abstract of the paper titles to get text documents for each of the Neurosynth papers. We then use a deep learning-based text embedding technique called Glove embeddings^38^ to convert these documents from Neurosynth and UK-Biobank to 300-dimensional vectors. This gives us two matrices: 1) 4926 × 300 dimensional matrix for UK-Biobank phenotypes and 2) 14371 × 300 matrix for Neurosynth papers. We then compute pairwise distances for each pair of phenotype and Neurosynth paper using inverse of cosine distance to obtain a 4926 × 14371 dimensional text similarity matrix.

### Brain and text alignment

Once the two similarity matrices were constructed, each giving an all-to-all comparison between Neurosynth papers and Biobank variables, one in text space and the other in brain space, with cell (*i, j*) of each matrix representing the same phenotype *i* and Neurosynth paper *j* pair, we computed an alignment score between the two matrices.

The alignment score was computed by flattening the two matrices into 7,246,146 dimensional vectors. Once we have these two vectors, we compute pair-wise Pearson correlation coefficient to give us an alignment score. This alignment score represents the degree of congruency or agreement between the two spaces, meaning if a paper-variable pair have highly similar activation maps, so that pair also has highly similar text descriptions.

## Notes

https://www.neuropredictome.com

### Competing Interest Statement

The authors have declared no competing interest.

